# Dude, Everyone Wants Pattern Analysis Tools (DEWPAT): Tools for measuring visual pattern diversity from digital images

**DOI:** 10.1101/2025.01.29.635361

**Authors:** Jillian A Sanderson, Tristan Aumentado-Armstrong, Charles-Olivier Dufresne-Camaro, D Luke Mahler

## Abstract

1. Exploring the diversity and function of complex colour patterns is a fundamental interest in ecology and evolutionary biology, but progress on many questions is limited by our ability to quantify diverse visual patterns. We address this problem by introducing Dude, Everyone Wants Colour Pattern Analysis Tools (`DEWPAT`), a Python package for characterizing multidimensional pattern complexity.
2. `DEWPAT` is a flexible framework designed to extract a diversity of components of visual pattern complexity from standard RGB and multispectral images, including entropy (information content), average gradient magnitude (edge content), high frequency content (detail granularity), generalized variance (heterogeneity), and patch dissimilarity. `DEWPAT` offers optional image transformation functionality, including blurring to model receiver acuity and segmentation to reduce noise.
3. Functions in this package return both quantitative measurements and graphical representations of color and pattern diversity. We demonstrate `DEWPAT`’s key functions and applications with three empirical examples (longhorn beetles, anole lizards, and flowers).
4. `DEWPAT` has the potential to quantitatively characterize previously intractable pattern phenotypes in ways that make their features available for biological analysis.

## 1. Introduction

Many species that communicate visually possess traits with elaborate and complex colour patterns (Pérez i de Lanuza & Font, 2016; Dalrymple et al., 2020), contributing to much of the phenotypic diversity we see in nature. Visual signals vary both within and among species, ranging from uniform colour patches to multi-colour designs with intricate spatial arrangements. Complexity is an emergent feature of patterns that can serve several important communication functions, including the exaggeration of sexual signals (Chen et al., 2012), augmentation of signal information content (Driessens et al., 2015), and modification of the efficacy of signal transmission, with complexity potentially reducing conspicuousness (e.g., camouflage; Chiao et al., 2009) or elevating it (e.g., social communication; Galván, 2008).

Interest in visual pattern complexity has led to an explosion of techniques to quantify the spectral and spatial components of patterns in digital images. For example, recent software can characterize the spatial arrangement of pattern elements and analyze similarity between images (Stoddard et al., 2014; Van Belleghem et al., 2017; Chan et al., 2019). Further, there are now techniques that combine spectrometer-collected colour information and pattern geometry, such as pattern contrast, adjacency, and boundary strength analyses (Endler & Mielke, 2005; Endler, 2012; Endler et al., 2018; Maia et al., 2019). More recently, there has been a focus on quantifying colour and pattern in a way that more faithfully reflects receiver perception. For example, new software creates image models that reflect the visual acuity (Caves & Johnsen, 2017) and colour vision (Troscianko & Stevens, 2015; Gawryszewski, 2018; van den Berg et al., 2019) of organisms. Finally, there has been an emerging interest in quantifying visual pattern complexity, with researchers developing methods to measure particular dimensions of complexity (e.g., Chen et al., 2012; Ligon et al., 2018).

Despite these recent innovations, tools for quantifying diverse aspects of visual signals remain fragmented. For example, many methods analyze spectral data without incorporating spatial information, and many pattern-focused methods measure and compare individual pattern elements rather than whole patterns, limiting their ability to quantify pattern complexity. Methods that do characterize whole colour patterns often focus on a single pattern dimension, failing to capture the complementary ways different patterns may be complex, which is important if complexity is a target of selection. Further, many pattern analysis methods obligatorily simplify images into discrete colour clusters, losing potentially useful information, especially for patterns featuring continuous colour gradients. Recent software advances have begun to combine spatial and spectral information, but these require knowledge of organisms’ visual systems, limiting their utility for exploratory or large-scale comparative studies.

More generally, a specific focus on colour pattern complexity is lacking in recent tools, potentially due to the difficulty of defining what makes a pattern complex. Visual complexity can be achieved in different ways, depending on the number and distribution of colour and pattern elements (Hebets & Papaj, 2005). Unidimensional measures of complexity constrain our ability to study the ecology and evolution of complex colour patterns. For example, a given complexity measure may capture pattern variation in one organism but not in another that varies along a different axis of pattern diversity. Additionally, within organisms, pattern differences along one axis could evolve independently or congruently with those along a different axis. The field of computer vision offers techniques to measure differentiation in complexity across different dimensions, but these have been underutilized in biology.

Here we introduce Dude, Everyone Wants Pattern Analysis Tools (`DEWPAT`), a Python package (Python v3.7; Van Rossum & Drake, 2009) for measuring complex colour patterns from digital images. `DEWPAT` combines new and existing image analysis tools into an easy-to-use framework. Developed from our own research on biological systems with diverse, multidimensional patterns, `DEWPAT` utilizes computer vision techniques to extract information about pattern complexity and colour distributions from images. `DEWPAT` integrates a variety of pre-processing, input, analysis, and output tools, making it versatile for breadth of applications. In this paper we describe `DEWPAT`’s pattern analysis tools and demonstrate its use with three worked examples.

## 2. Methods

**Figure 1.**
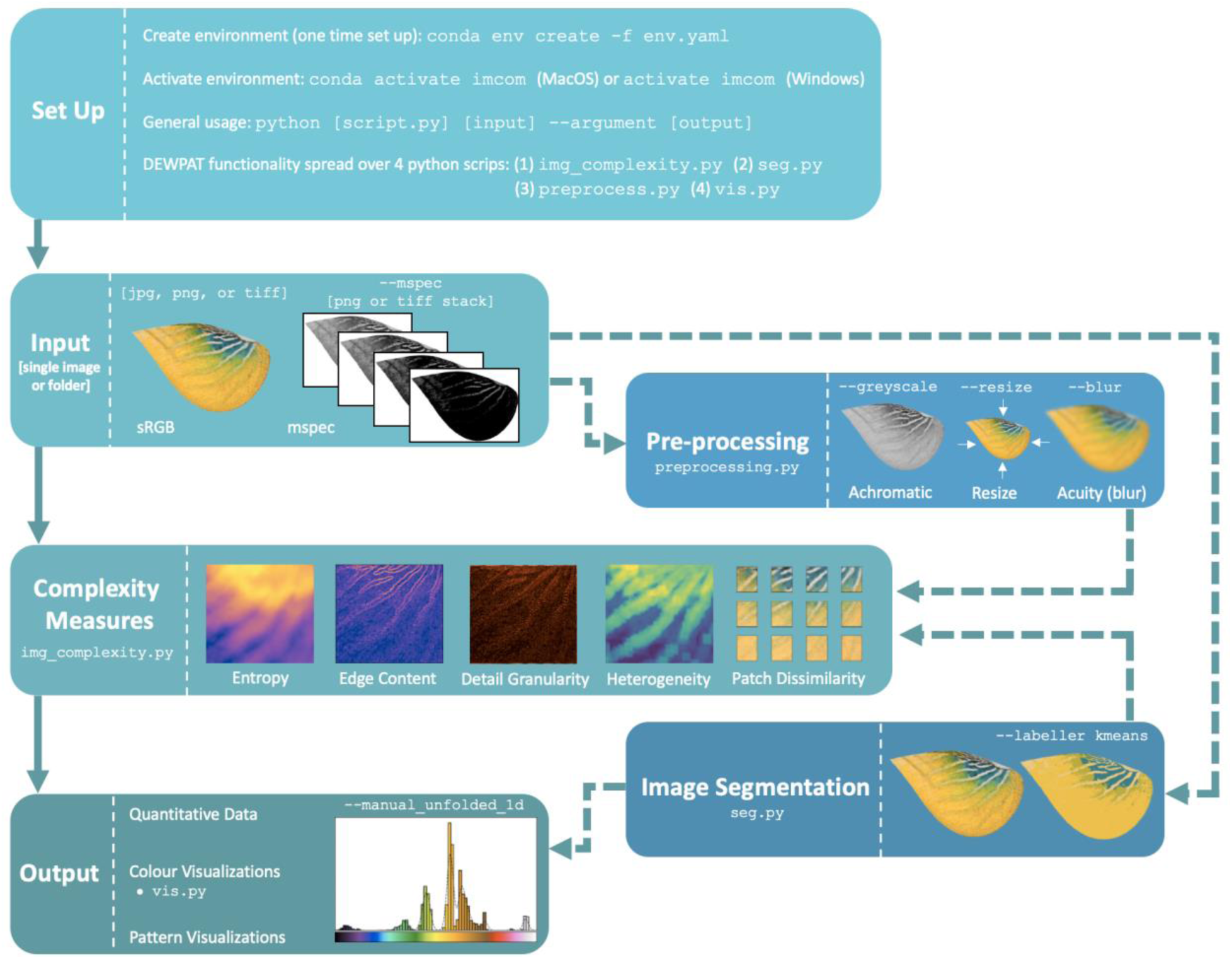
Overview of `DEWPAT` workflow. Solid lines indicate a basic analysis pipeline, while dashed lines indicate alternative pipelines that incorporate optional additional steps.

### 2.1. Input

`DEWPAT` can analyze standard RGB images or multispectral image stacks (by passing `--mspec`), where each image represents a different color channel for a single subject (e.g., including a UV wavelength channel; Troscianko & Stevens, 2015). See the Supporting Information for discussion of `DEWPAT` input requirements, including examples.

### 2.2. Complexity Measures

#### 2.2.1 Entropy (Information Content)

Entropy is a property of a distribution that measures information or uncertainty (Shannon & Weaver, 1949). There is interest in applying entropy to biological studies of signal complexity (reviewed in Gao et al., 2012). For example, entropy has been used to study information in anole displays (Rand & Williams, 1970; Nelson & Ord, 2022), to quantify alarm call complexity in chickadees (Freeberg, 2006), and to analyze multimodal signal complexity in birds-of-paradise (Ligon et al., 2018). `DEWPAT` can extract two types of entropy from images: discrete and differential. The discrete entropy measure assumes pixel values are numbers in a discrete set (here, a colour channel) and therefore computes entropy per colour channel and then averages these scores yielding a total discrete entropy value for the pattern (Figure 2B). By contrast, the differential entropy measure doesn’t separate the colour channels; instead, the image is treated as a continuous distribution of pixels, in which each pixel is represented by an N-dimensional vector and thus retains its individual meaning (e.g., red pixels are treated as more similar to orange pixels than to blue pixels). Entropy is computed over the distribution of all pixel vectors. Discrete and differential entropy can be computed across an entire image `(--discrete_global_shannon, (--diff_shannon_entropy)` or within-image patches that are then averaged (`--discrete_local_shannon, --diff_shannon_entropy_patches)`.

**Figure 2.**
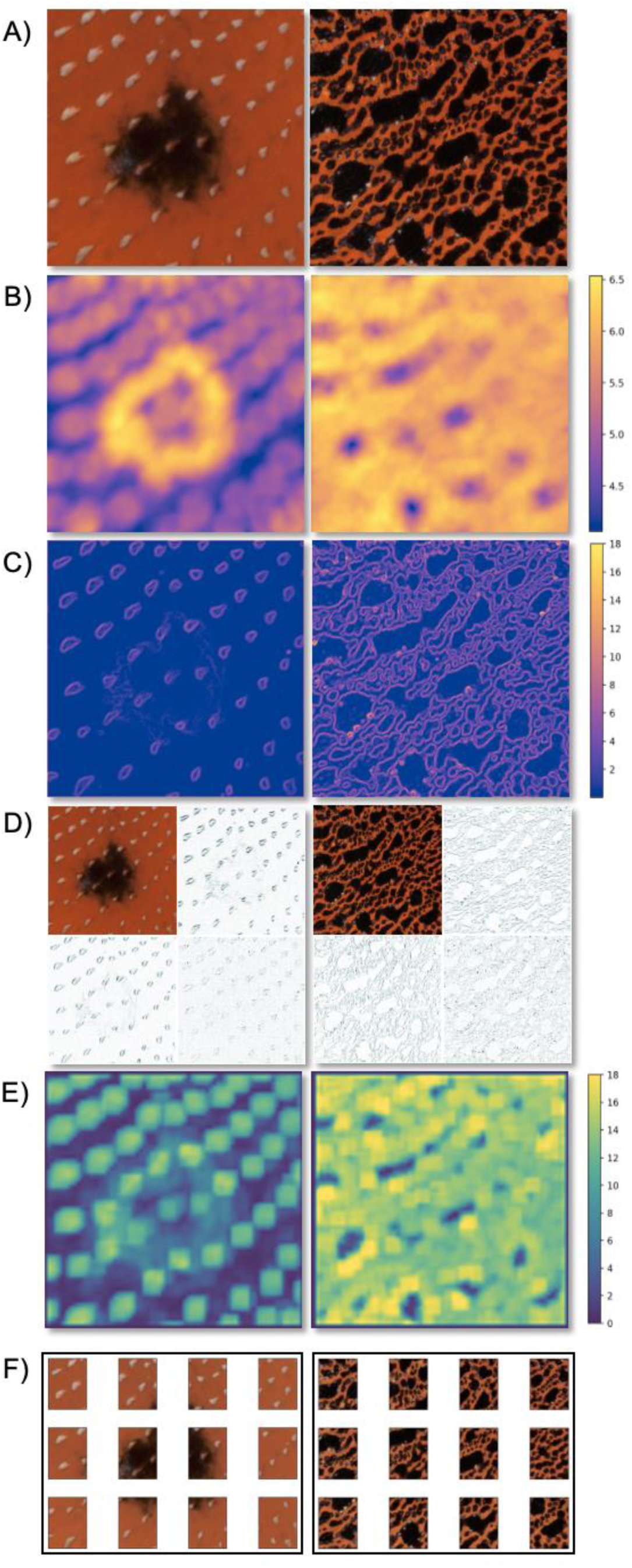
Visual output of different pattern complexity dimensions for selected image regions of two male *Anolis* lizard dewlaps (*Anolis lyra on left; A. antioquiae on right*). From top to bottom: original image samples (imported with `python img_complexity.py image_folder`), patch-wise discrete entropy `(--show_local_ents)`, average gradient magnitude `(--show_grad_mag)`, discrete wavelet transform `(--show_dwt)`, local heterogeneity `(--show_local_covars)`, and patch dissimilarity `(--show_pw_mnt_ptchs)`.

#### 2.2.2 Average Gradient Magnitude (Edge Content)

Edges provide boundaries between distinct adjacent pattern elements and are key to the perception of whole colour patterns, with the number and intensity of edges contributing to overall visual complexity (Endler, 2012; Endler et al., 2018). We extract edge magnitude from images by computing an image gradient consisting of the derivatives (local rates of change) of the pixel values over space. For each channel, we compute a vector indicating the direction and magnitude of this change at each image coordinate. We then take the average vector magnitude across channels as a measure of complexity `(--grad_mag`; Figure 2C).

#### 2.2.3 High Frequency Content (Detail Granularity)

Granularity, or high frequency content, refers to the level of signal detail and is increasingly used to characterize colour patterns in nature, especially for studies of camouflage and disruptive colouration (Godfrey et al., 1987; Chiao et al., 2009; Troscianko & Stevens, 2015). We can study granularity using spatial frequency analyses, which can convert signals from a spatial domain to the frequency domain, where information is represented by a series of waves with different spatial frequencies. These methods are based on animal neurophysiological spatial scene processing (Meese, 2002; Stoddard & Osorio, 2019). The transform coefficients describe the relationship between original and transformed signals, with high frequency coefficients representing signal regions containing high levels of detail (changes over short spatial distances) and low frequency coefficients representing signal regions containing low levels of detail (changes over large spatial distances).

We implement two spatial frequency analyses: Fourier transforms and discrete wavelet transforms, which use different base functions convert a signal into the frequency domain. Taking the Fourier transform of an image involves decomposing it into infinitely long sinusoidal waves with differing frequencies. We quantify image high frequency content (i.e., detail) by computing the weighted average of the Fourier coefficient values, with weights based on the associated frequency value `(--weighted_fourier)`. In contrast, wavelet transforms decompose a signal into a set of wavelets (small, localized waves). Wavelet coefficients are filtered into approximation (low frequency) and detail (high frequency) coefficients. We perform a 4-level discrete wavelet transform using the Haar mother wavelet (Haar, 1910) and compute the sum of the detail coefficients as our measure of high frequency content `(--wavelet_details)`. For simplicity, Figure 2D shows the original image and the resulting horizontal, vertical, and diagonal detail images after a single level of high-frequency filtering (see Supporting Information for additional details and images).

#### 2.2.4 Generalized Variance (Heterogeneity)

Heterogeneity can be defined as the quality of consisting of dissimilar elements. One method to characterize the heterogeneity of an image is to compute the covariances between pixel values and then calculate the covariance matrix determinant. This determinant, (aka the generalized variance; Wilks, 1932), is a scalar measure of the multidimensional spread of a set of observations (Bagai, 1965). We quantify generalized variance for images at two scales, local (within-patch) and global (between-patch). For both methods we start by dividing the image into small patches (default: 20×20 pixels). At the local scale, we compute the generalized variance of each patch and then take average patch score as a measure of complexity (`--local_covars;` Figure 2E). At the global scale, we treat each patch as a single high-dimensional vector, (dimension = number of channels x number of pixels per patch), compute the covariance across those vectors, and take the generalized variance of the between-patch covariance matrix `(--global_patch_covar)`.

#### 2.2.5 Pairwise Distance Measures (Patch Dissimilarity)

Pixel heterogeneity within and among small patches may fail to capture larger-scale structural complexity in patterns (e.g., if patches are much smaller than pattern elements). We have implemented a suite of patch dissimilarity measures to account for such larger pattern elements. For these methods, we first divide the image into a number of large, non-overlapping patches (default: 12; Figure 2F). We then characterize the pattern of each patch using moments, with the first moment being the mean value of the pixels in the patch (i.e., average colour) and the second moment being the covariance of pixels in that patch. To characterize coarse-scale pattern information, we can calculate the average distance between patches using the first moment as a measure of colour distribution, or both moments as measures of colour and pattern distribution. Since some vector space metrics (e.g., Euclidean distance) poorly reflect biological visual perception (Zhang et al., 2018), we utilize several distributional divergences instead, which are more robust to perceptually irrelevant perturbations, such as small translations (see the Supporting Information for details).

### 2.3. Additional Functionality

Investigators may wish to transform images for denoising purposes or to account for receiver perception prior to analysis. We have implemented the AcuityView algorithm (Caves & Johnsen, 2017) in `preprocess.py` to blur sRGB and mspec images, modeling how patterns are perceived based on visual acuity and distance. `DEWPAT` also includes numerous colour segmentation methods for sRGB images in `seg.py`. The default method is k-means clustering (Lloyd, 1982) where k can be manually specified (e.g., `--kmeans_k 4`) or automatically detected; users may also choose to cluster pixels by mean `(--write_mean_segs)`, median `(--write_median_segs)`, or modal `(--write_mode_segs)` value. We provide a detailed explanation of all segmentation options (including examples) in the Supporting Information.

### 2.4. Output

`DEWPAT’s` main pattern analysis script, `img_complexity.py,` generates scores for all complexity measures for each image. `seg.py` provides quantitative output containing colour codes (in three colour spaces: RGB, HSV, and CIELAB) and the pixel frequency of each cluster for each image. For RGB images, `DEWPAT` can output numerous visual representations of pattern complexity, colour segmentation, and colour distribution (e.g., a 1D colour histogram; Figure 3). For segmented images, `DEWPAT` can also return a colour transition matrix based on Endler’s adjacency analysis (see Endler, 2012 for detailed description). Additional output options are described in the Supporting Information.

**Figure 3.**
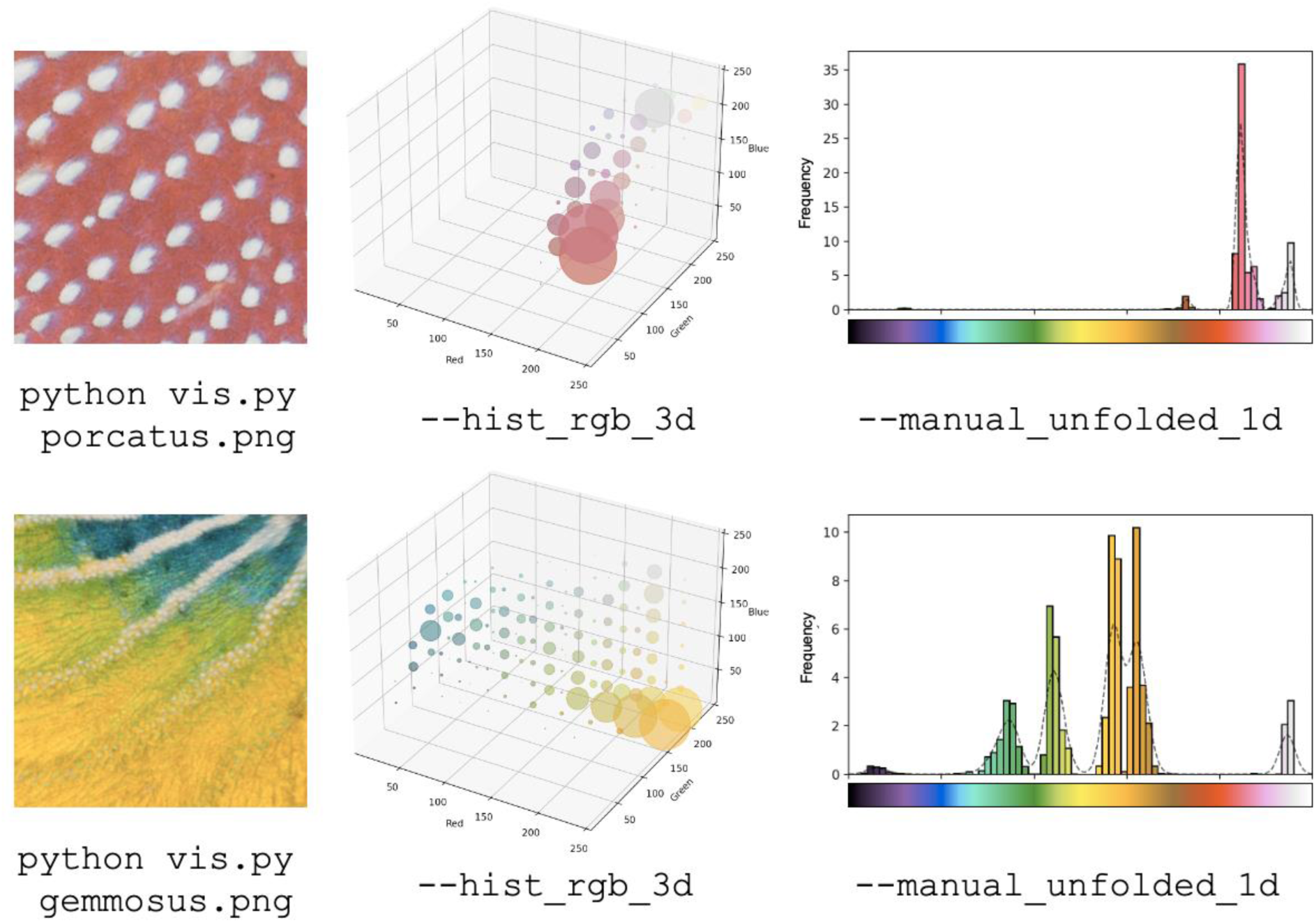
Example colour visualization and code. Left: selected regions of male *Anolis* lizard dewlaps (*Anolis porcatus* top; *A. gemmosus* bottom). Middle: 3D pixel frequency plots, where point size corresponds to pixel frequency. Right: 1D pixel histogram (see Supporting Information for details).

## 3. Examples

### Replication Statement

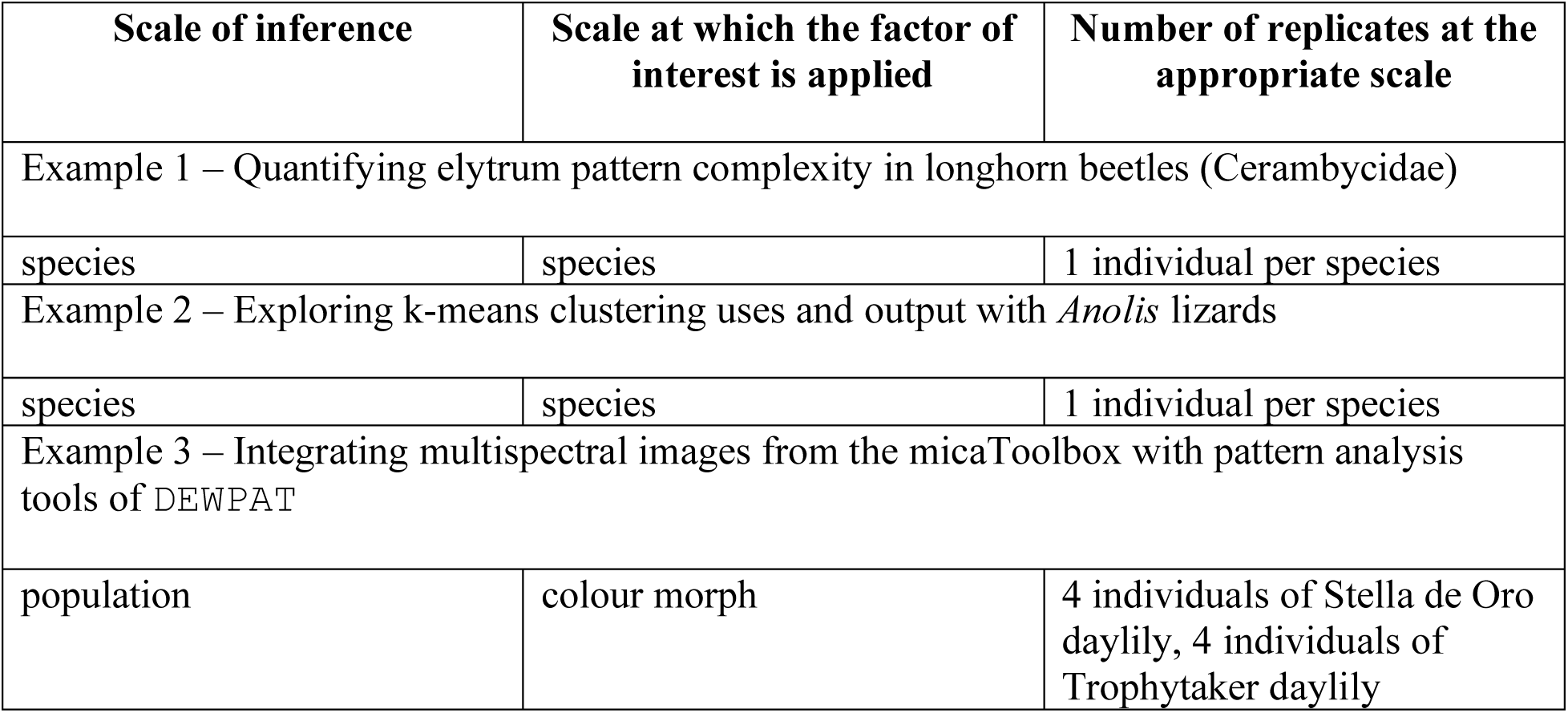

| Scale of inference | Scale at which the factor of interest is applied | Number of replicates at the appropriate scale |
| --- | --- | --- |
| Example 1 – Quantifying elytrum pattern complexity in longhorn beetles (Cerambycidae) |  |  |
| species | species | 1 individual per species |
| Example 2 – Exploring k-means clustering uses and output with *Anolis* lizards |  |  |
| species | species | 1 individual per species |
| Example 3 – Integrating multispectral images from the micaToolbox with pattern analysis tools of DEWPAT |  |  |
| population | colour morph | 4 individuals of Stella de Oro daylily, 4 individuals of Trophytaker daylily |

### 3.1. Example 1 – Quantifying elytrum pattern complexity in longhorn beetles (Cerambycidae)

We first demonstrate `DEWPAT`’s utility by measuring colour pattern complexity in the elytra of longhorn beetles (Cerambycidae). We compiled elytrum images of 30 species from the family Lamiinae from an online gallery (Roguet, 2022), and used `DEWPAT` to extract four features of pattern complexity from these images: patchwise differential entropy, average gradient magnitude, global patch covariance, and Bhattacharyya divergence (i.e., patch dissimilarity).

`python img_complexity.py beetle_elytra`
`--diff_shannon_entropy_patches --grad_mag`
`--global_patch_covar --pwg_bhattacharyya_div`
`>beetle_complexity.csv`

To reduce the dimensionality of this dataset, we analyzed the complexity scores with a principal component analysis (PCA) using the `prcomp()` function in `Rv4.0` (R Development Core Team). The first two principal components explain 87% of the pattern variation in the beetle elytra. PC1 is most strongly correlated with patterns containing many small details, (i.e., fine-scale complexity), and PC2 is strongly correlated with patterns containing few, bold markings (i.e., coarse-scale complexity) (Figure 4). By providing a variety of pattern analysis tools, `DEWPAT` can meaningfully distinguish signals where patterns not only range from simple to complex, but where complex patterns diverge on different complexity axes.

**Figure 4.**
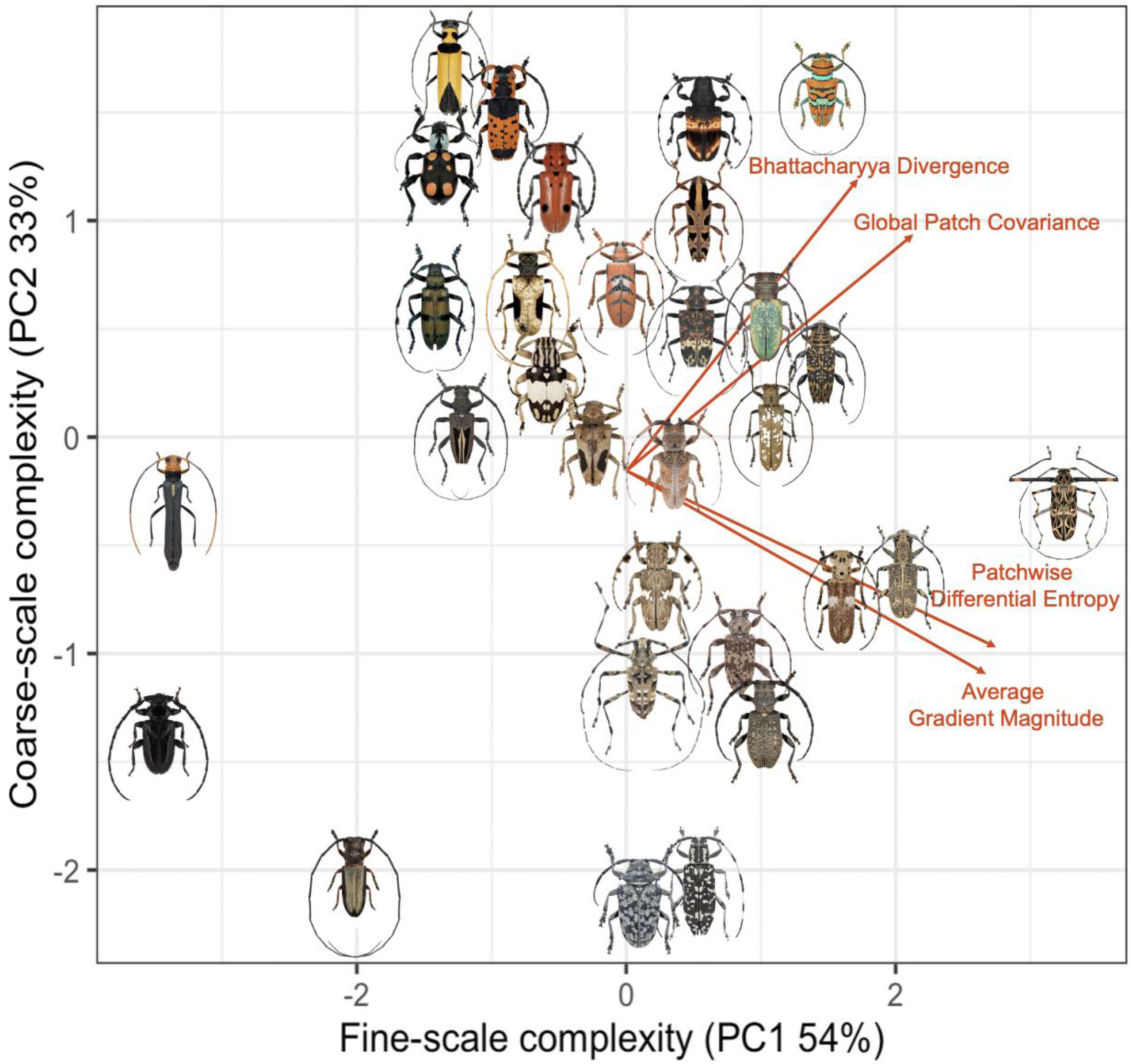
Biplot of the first two principal components from a PCA of four `DEWPAT` complexity measures. Red lines indicate the loadings of the original variables relative to PCs 1 and 2. Some beetle images were jittered by up to one body width to reduce overlap.

### 3.2. Example 2 – Exploring k-means clustering with Anolis lizards

Our next example demonstrates `DEWPAT’s` colour segmentation functionality using the colourful and diversely-patterned dewlaps (throat fans) of *Anolis* lizards. We used `DEWPAT`’s auto-segmentation feature to segment images of adult male dewlaps for six *Anolis* species into k-clusters, coloured by median colour. We output a file that, for each image, reports the colour value in LAB space of each detected colour cluster and the frequency of pixels in each.

`python seg.py anole_dewlaps`
`--labeller kmeans`
`--write_median_segs`
`--median_seg_output_dir anole_segs`
`--seg_median_stats_output_file anole_colour_stats.csv`

We used this output to determine the two most dominant dewlap colours for each species. Finally, we calculated the Euclidean distance between these two colours as a measure of within-dewlap colour contrast (Ortiz-Jaramillo et al., 2016; Figure 5), a signal attribute known to increase conspicuousness (Pérez i de Lanuza & Font, 2016).

**Figure 5.**
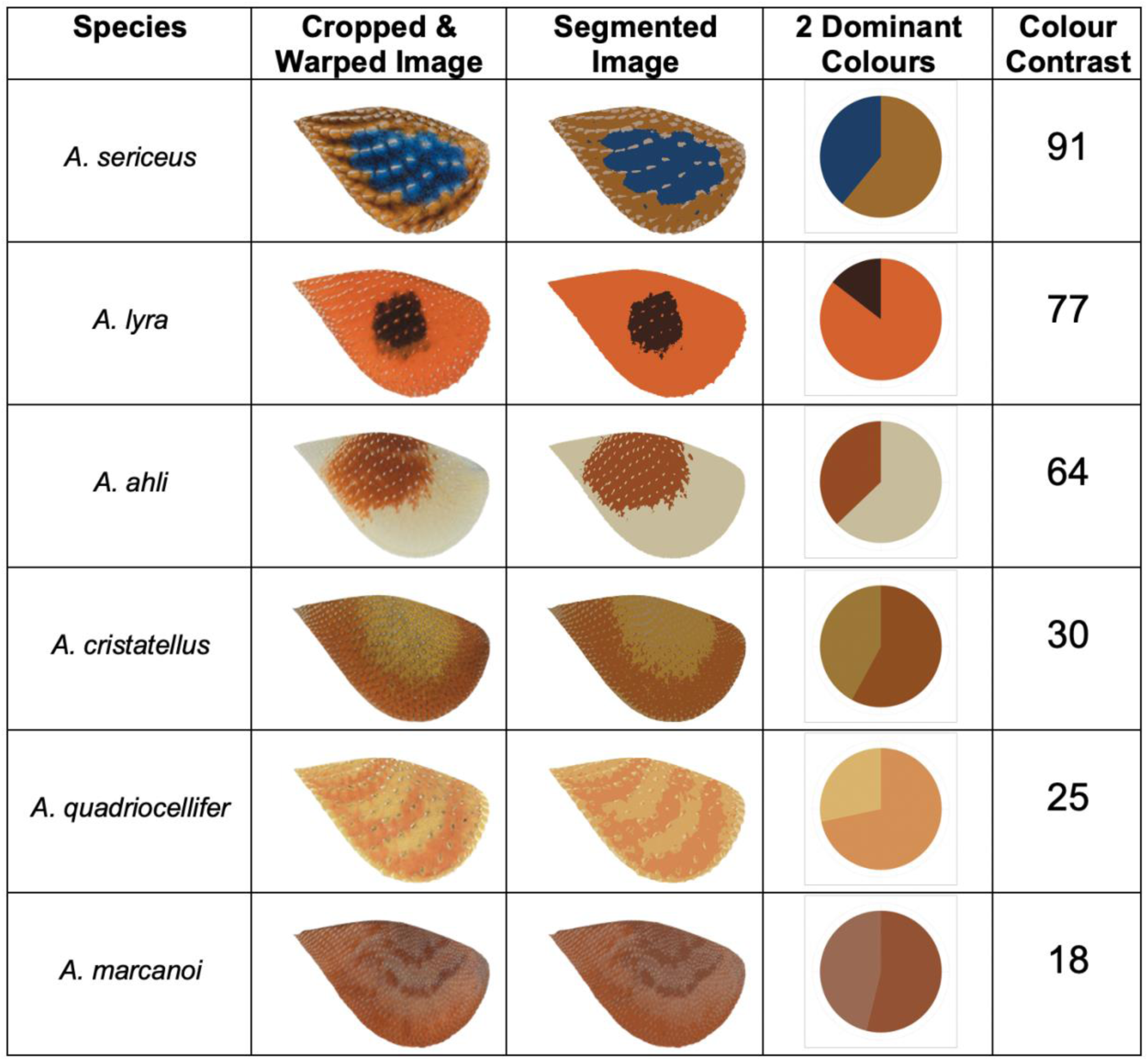
Dewlap colour contrast of 6 *Anolis* species sorted from highest to lowest colour contrast.

### 3.3. Example 3 – Integrating multispectral images from the micaToolbox with DEWPAT pattern analysis tools

Recent advances in visual modeling have made it possible to account for the visual system and perceptual capabilities of organisms, allowing us to better understand how a colour pattern may appear to various animal viewers (Caves & Johnsen, 2017, Troscianko & Stevens, 2015, van der Berg et al., 2020). Here we compare visual system-corrected and non-corrected images of flower petals from two daylily (*Hemerocallis lilioasphodelus*) colour morphs (Figure 6) to demonstrate `DEWPAT`’s ability to handle multiple image types, and how different visual models can lead to different conclusions about patterns.

**Figure 6.**
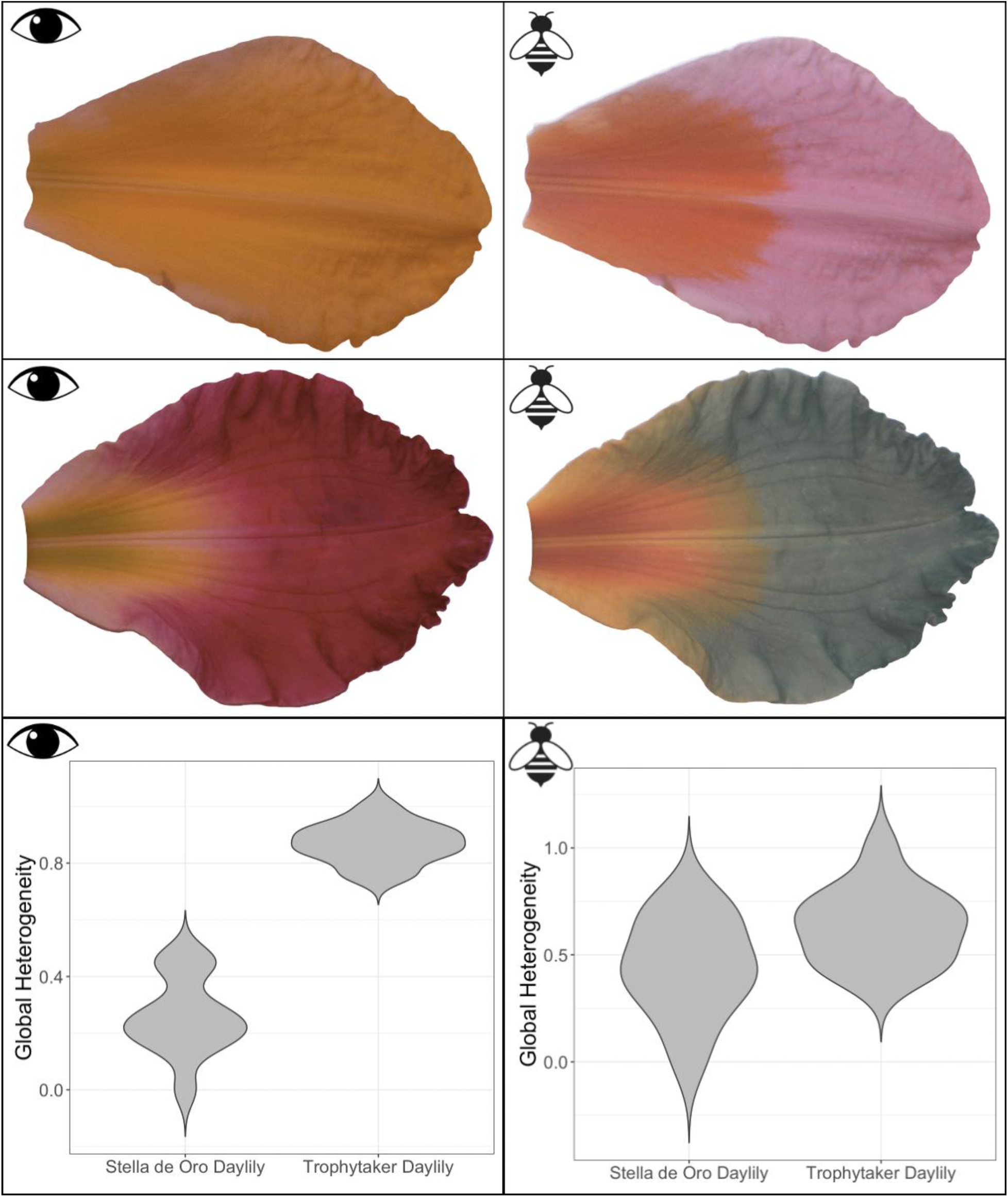
Comparison of flower petals from two daylily horticultural varieties (top row: Stella de Oro, middle row: Trophytaker) as seen from the human visual system (left) and honeybee visual system (right). Violin plots show colour pattern heterogeneity extracted from flower petal images representing the two visual systems. These daylily varieties differ more in appearance to humans than they do to a honeybee. Note that the false coloured honeybee corrected images were created with the `micaToolbox` plugin (Troscianko & Stevens, 2015) in `ImageJ` (Schneider et al., 2012).

We used two sets of flower petal images: (1) RGB images representative of typical digital camera images (hereafter: human-vis images) and (2) multispectral image stacks corrected to reflect the perceptual space of the honeybee (*Apis mellifera*) using the `micaToolbox` (Troscianko & Stevens, 2015) in `ImageJ` (Schneider et al., 2012) (honeybee-vis images). We first used `DEWPAT` to measure global heterogeneity from all images.

`python img_complexity.py flower_ex_vis_flowers`
`--global_patch_covar >flower_vis_complexity.csv`

`python img_complexity.py --mspec flower_ex_bee_flowers`
`--global_patch_covar >flower_bee_complexity.csv`

We then used ANOVA to compare the colour patterns of the two flower morphs in both honeybee-vis and human-vis spaces in `R v4.0` (R Development Core Team). For the human-vis images, there is a large difference in pattern heterogeneity, and the Trophytaker daylily is more heterogeneous than the Stella de Oro daylily (Figure 6; F = 186.2, p < 0.001). Under the honeybee visual model, the pattern difference between morphs is much less pronounced (F = 5.3, p = 0.032), indicating that the flowers appear more similar to a biologically relevant receiver (Figure 6). By accommodating multispectral images, `DEWPAT` can harness the powerful visual modeling capabilities of the `micaToolbox` (Troscianko & Stevens, 2015), allowing researchers to analyze colour patterns from visual system-corrected images.

## Conclusion

`DEWPAT` addresses the challenges of studying multidimensional patterns by providing a new toolkit for quantifying and illustrating visual pattern complexity from digital images. We integrate both classic and novel pattern-measurement tools with recently-developed techniques for representing diverse visual systems into a comprehensive framework. Ultimately, we hope `DEWPAT` will enable scientists to quantitatively characterize previously intractable phenotypes, paving the way for deeper insights into colour pattern function, ecology, and evolution.

## Notes

https://gitlab.com/taumen/image-complexity

### Competing Interest Statement

The authors have declared no competing interest.

