## Supporting Information for "Dude, Everyone Wants Pattern Analysis Tools (DEWPAT): Tools for measuring visual pattern diversity from digital images"

TABLE OF CONTENTS:

S1: GETTING STARTED WITH DEWPAT

S2: SUPPLEMENTAL DEWPAT INPUT REQUIREMENTS

(a) Single image or folder of images

(b) Image types and file formats

(c) Alpha mask

(d) Standardizing images

(e) Example workflow for size and shape standardization of images with irregularly shaped ROIs

S3: SUPPLEMENTAL COMPLEXITY MEASURE EQUATIONS

(a) Information Content/Entropy

(b) Edge Content/Average Gradient Magnitude

(c) Detail Granularity/High Frequency Content

(d) Heterogeneity/Generalized Variance

(e) Patch Dissimilarity/Pairwise Distance Measures

S4: SUPPLEMENTAL DESCRIPTION OF DISCRETE WAVELET TRANSFORM (DWT) OUTPUT IMAGE

S5: SUPPLEMENTAL IMAGE SEGMENTATION DESCRIPTION AND EXAMPLE

S6: SUPPLEMENTAL DATA OUTPUT AND VISUALIZATION OPTIONS

S6: SUPPLEMENTAL REFERENCES

### S1: GETTING STARTED WITH DEWPAT

#### 1. Download Python

Users can run python directly through the command line. For users unfamiliar with python, we suggest using an Integrated Development Environment (IDE) (e.g., PyCharm) to run scripts.

#### 2. Clone/download the DEWPAT repository from the GitHub page

DEWPAT currently exists in the GitHub page https://github.com/jillsanderson/DEWPAT/ as a series of Python scripts divided between two branches. Note that both branches should technically contain all of the scripts, but for best and most up to date functionality users should use the script from the branch designated below.

*master branch*

- img_complexity.py: main script that includes functionality to compute and visualize complexity across both sRGB and multispectral images. Note that if using mspec images, need to run `--mspec` along with other arguments
- vis.py: includes visualization options for sRGB images (including 1D colour histogram)

*dev branch*

- seg.py: includes functionality to segment, visualize, and output colour statistics for sRGB images
- preprocess.py: includes functionality to blur both sRGB and multispectral images to model visual acuity and viewing distance using AcuityView (Caves & Johnsen, 2017).

Note that everything is run out of the repository folder (i.e., where the python scripts exist). Make sure to add the images for analyzes to this folder.

##### 3. Set up python

DEWPAT scripts require: `scikit-image, numpy, matplotlib, scikit-learn, and scipy.` An easy way to obtain them is via the conda environment specified in the env.yaml file included when you clone the package from the GitHub repository.

Create the environment by running: `conda env create -f env.yaml` (this only has to do this once, the first time DEWPAT is set up)

To activate the environment once it's created run: `conda activate imcom` (MacOS) or `activate imcom` (Windows)

##### 4. Run scripts

The general usage is as follows:

```
python [script_name.py] [file.extension or input folder] --usage_specific_argument [output.csv]
```

For example, to calculate the detail coefficients for a png image called example_image using a discrete wavelet transform, display a visualization of the transform, and export results to a .csv file, you would run:

```
python img_complexity.py example_image.png --wavelet_details --show_dwt >image_wavelet_output.csv
```

To print a detailed help page for each script run: `python [script_name.py] --help`. ***We recommend doing this for each script to get a full breakdown of all commands and script functionality.***

Data necessary to replicate examples and figures from the main text (i.e., images, .csv files, and code) can be found in GitHub in the DEWPAT_manuscript_examples folder.

We note that if users wish to run scripts using default algorithm parameters (e.g., patch sizes, bin numbers, etc.) then they can simply run commands in the terminal. If users wish to change default parameters, they first need to find and edit the parameters in the scripts themselves. See the main text for details about default parameters.

See README.md files on GitHub for more information including examples.

#### S2: SUPPLEMENTAL DEWPAT INPUT REQUIREMENTS

##### (a) Single image or folder of images

DEWPAT operates on digital images, taking a single image or a folder of images as input. If inputting a folder of images, all images should have the same dimensions (e.g., all 500x100 pixels) and be in the same file format (e.g., all PNGs).

##### (b) Image types and file formats

DEWPAT can analyze standard red, green, blue (sRGB) images or multispectral (mspec) image stacks. RGB is a colour space consisting of the colours red, green, and blue and is designed to approximately align with human photoreceptors. Original (i.e., raw) images are roughly linearly related to scene irradiance (so-called linear RGB) and standard cameras then use a camera pipeline (or image signal processor) to transform it to sRGB (see Karaimer and Brown, 2016 for an overview). An sRGB image is thus an image with three colour channels (red, green, and blue) captured and expressed as pixel values within a single image. sRGB is the most widely used colour space but as it comes with some assumptions about the receiver (e.g., human viewer with no colour vision deficiencies), we also support multispectral images. For RGB images, files can be in JPEG, PNG, or TIFF format.

A multispectral image stack is collection of normalized, aligned images, each captured at a particular wavelength band. Multispectral image stacks allow image channels with often-excluded wavelengths to be represented (e.g., via a UV channel), and can be converted to cone catch quanta of different visual systems (Troscianko and Stevens, 2015). In an mspec image stack, images appear in greyscale because the pixel values of each image represent the amount of light stimulating the photoreceptor being modeled by that image. For mspec image stacks, files can be TIFF stacks (i.e., one file that contains n images, depending on number of channels) or sets of PNG images (i.e., n files, each a single image representing a single channel). Note that if inputting sets of PNGs, the user must put all the images loose in the main folder (not nested within their own folder) and include a .csv file indicating which images correspond to which set (Figure S1). See Table S1 for an overview of the differences between RGB and mspec images.

| | A | B | C |
|---|---|---|---|
| 1 | ids | stacknum | filename |
| 2 | 0 | 0 | specimen_0016_1.png |
| 3 | 1 | 0 | specimen_0016_2.png |
| 4 | 2 | 0 | specimen_0016_3.png |
| 5 | 3 | 0 | specimen_0016_4.png |
| 6 | 4 | 1 | specimen_0017_1.png |
| 7 | 5 | 1 | specimen_0017_2.png |
| 8 | 6 | 1 | specimen_0017_3.png |
| 9 | 7 | 1 | specimen_0017_4.png |
| 10 | 8 | 2 | specimen_0030_1.png |
| 11 | 9 | 2 | specimen_0030_2.png |
| 12 | 10 | 2 | specimen_0030_3.png |
| 13 | 11 | 2 | specimen_0030_4.png |
| 14 | 12 | 3 | specimen_0036_1.png |
| 15 | 13 | 3 | specimen_0036_2.png |
| 16 | 14 | 3 | specimen_0036_3.png |
| 17 | 15 | 3 | specimen_0036_4.png |

**Figure S1.** Example format of .csv file that indicates which images correspond to which PNG set. Files must follow this format to be recognized by DEWPAT.

**(c) Alpha mask**

By default, the entire input image will be analyzed by DEWPAT and the package does not currently have the option to specify regions of interest (ROIs). However, most measures are able to account for the presence of an alpha channel mask, which allows the user to run the measures on a pre-specified ROI rather than the entire image. An alpha mask is a separate image channel in PNG and TIFF images that accounts for image transparency. For RGB images, a simple way to include an alpha mask is to first remove the background or any region of the image you do not want analyzed (e.g., using Batch-Mask; Curlis et al. 2022 or a photo editing software such as Adobe Photoshop™, and then save the image as a PNG. Typically, PNG images have an alpha channel (this is why PNG images can have transparent backgrounds) but JPEGs do not. For mspec PNG sets, an alpha mask can be created in the same manner as a standard RGB image, and for mspec TIFF stacks, an alpha mask can be created by setting the background pixels to NaN using ImageJ™ or similar software.

**(d) Standardizing images**

For most consistent results, we recommend preparing images so that they exhibit standardized size, shape, and lighting conditions before inputting them. Colour calibration can be accomplished through the use of colour and luminance standards during image capture (e.g., an X-Rite ColorChecker Passport), followed by calibration to standardized colour profiles during image pre-processing. When analyzing rectangular sections of images, size and shape can be standardized by cropping images to be the same dimensions (which is recommended when inputting a folder of images). When analyzing irregular shaped patterns or body regions, size and shape can be standardized by performing image registration (i.e., by applying landmarks and performing a Procrustes superimposition of images).

**(e) Example workflow for size and shape standardization of images with irregularly shaped ROIs**

- In Adobe Lightroom™, crop each image to focus only on the ROI using a 1x1 aspect ratio
- Resize all images to the dimensions of the smallest image to reduce image enlargement distortion
- Use these cropped, re-sized images to create a TPS file using TPS Util™ (Rohlf, 2004)
- Landmark around the ROI perimeter in each image with sliding semi-landmarks using TPS Dig™ (Rohlf, 2004)
- Use TPS Super™ (Rohlf, 2004) to perform a Procrustes superimposition, warping all images to the mean consensus shape (or to a target shape)
- Load warped images as layers into Adobe Photoshop™ and then remove background by creating a clipping mask
- Save layers to PNG files and make sure to check the "include transparency box" to create the alpha mask

**Table S1.** General overview of the differences between sRGB and mspec images.

| | **RGB** | **mspec** |
|---|---|---|
| **Description** | Standard red, green, blue (sRGB) is the current default colour space for computers. Most images (other than RAW) are in the sRGB format by default. | Multispectral (mspec) images are stacks of images taken at different wavelengths. These start as RAW images and are later saved into TIFF stacks or sets of PNG images. |
| **Pixel values** | A 3D vector containing values for the red channel, the green channel, and the blue channel (together referred to as an ‘RGB triplet’). Each value ranges from 0-255. Together, the combination of these values represents a colour. | A single reflectance value at a given wavelength. Values range from 0-1. Images are displayed in greyscale, with darker pixels having values closer to 0 and lighter pixels having values closer to 1. |
| **Number of images** | A single image. Although an RGB image is technically a multispectral image displaying three colour channels, each pixel contains all three of these values, thus allowing the use of one image rather than three separate images to display the colours. | Multi-image stack, where n = number of wavelengths being represented. As each pixel value contains only one reflectance value, mspec image stacks are comprised of as many images as there are represented wavelengths. |
| **Visual modeling** | Cannot be used for visual system modeling. | Can be used for visual system modeling. |
| **File formats** | JPEG, PNG, TIFF | TIFF stack, PNG set |
| **Alpha channel masking** | Remove background or any other portion of image you don’t want to analyze (i.e., using Photoshop™ or similar application). Only works with PNGs, which can have transparent backgrounds. | For TIFF stacks, set background or any other portion of image you don’t want to analyze to NaN (i.e., using imageJ or similar application).<br><br>For PNG sets, follow sRGB procedure. |

#### S3: SUPPLEMENTAL COMPLEXITY MEASURE EQUATIONS

We briefly provide a basic description of the measures and equations. See the code for precise computational details. General notation: image $I$ with channels $c \in C$, set of patches $P$ (per channel, $P_c$), set of pixels $V$ (per channel, $V_c$), and empirical covariance matrix $\hat{C}$.

##### (a) Entropy (Information Content)

Notation: $\mathcal{H}$ = entropy, p = probability of pixel value s (here, empirical frequency), $C$ = channel set, $V_c$ = pixels of channel c, d,v = “discrete” + “values”, d,p = “discrete” + “patch”.

###### *Pixel-wise Discrete Entropy*

Computes the discrete Shannon entropy over individual pixel values across the image:

$$\mathcal{H}_{d,v}(I) = \frac{-1}{|C|}\sum_{c\,\in\,C}\sum_{s \in V_c} p(s) log\, p(s)$$

`Python command: --discrete_global_shannon: calculate pixel-wise discrete entropy`

###### *Patch-wise Discrete Entropy*

Computes the mean discrete Shannon entropy within local image patches (overlapping disks centred at every pixel with a radius of 24 pixels by default). Notation: $\zeta$ = the space of discrete pixel values.

$$\mathcal{H}_{d,p}(I) = \frac{-1}{|C|}\sum_{c\,\in\,C}\frac{-1}{|P_c|}\sum_{\zeta \in P_c}\sum_{v \in \zeta} p(v)\, log\, p(v)$$

```
Python commands: --discrete_local_shannon: calculate patch-wise discrete entropy; --show_local_ents: display visualization
```

##### *Pixel-wise Differential Entropy*

Computes the differential Shannon entropy of the continuous-space vector-valued pixel distribution of the entire image. Notation: c stands for "continuous", and once again we use v to mean "pixel values")

$$\mathcal{H}_{c,v}(I) = -\int_V p(v)\, log\, p(v)\, dv$$

```
Python command: --diff_shannon_entropy: calculate pixel-wise differential entropy
```

##### *Patch-wise Differential Entropy*

Computes the differential entropy of the distribution of patches over the images (square patches of 20x20 pixels by default), where each multi-channel patch is unfolded into a single continuous vector. Notation: Integration is over space of patches P, where $\xi$ is a patch and p = probability of pixel value s.

$$\mathcal{H}_{c,p}(I) = -\int_P p(\xi)\, log\, p(\xi)\, d\xi$$

Note: the differential entropy measures utilize the Non-parametric Entropy Estimation Toolbox by Greg Ver Steeg (Steeg, 2000) and implement the approach in Kraskov et al. (2004). We also note that both entropy measures treat pixel values as independent across an image (or patch), and thus no do not quantify spatial structure.

```
Python command: --diff_shannon_entropy_patches: calculate patch-wise differential entropy
```

**(b) Average Gradient Magnitude (Edge Content)**

***Average Gradient Magnitude***

Computes the mean value of the per-channel gradient magnitude over the image using the following equation. Notation: ∇ = spatial gradient of the image.

$$\mathcal{G}(I) = \frac{1}{|C||V_c|}\sum_{c\,\in\,C}\sum_{s\in V_c}||\nabla I(s)||_2$$

```
Python commands: --grad_mag: calculate average gradient magnitude; --show_gradient_img: display visualization
```

**(c) High Frequency Content (Detail Granularity)**

***Frequency-Weighted Mean Fourier Coefficients***

Coefficients are computed as the weighted average of the Fourier coefficient values, with weights based on the associated frequency value, such that:

$$\mathcal{F}_\omega(I) = \frac{1}{Z_\gamma}\sum_{\psi_x,\psi_y\in\Psi}\gamma(\psi_x,\psi_y)\mathfrak{I}_F(\psi_x,\psi_y),$$

where $\Psi$ is the set of spatial frequencies (Fourier-space coordinates), $\gamma(x,y) = |x| + |y|$ is the Manhattan distance weighting function, and we denote

$$\mathfrak{I}_F = \frac{1}{|C|}\sum_{c\,\in\,C} log|\Im[c]||,$$

with $\Im[c]$ as the Fourier transform of the single-channel image *c*, *and* $Z_{\gamma} = \sum_{\psi_x,\psi_y \in \Psi} \gamma(\psi_x, \psi_y)$ is a normalization constant. Intuitively, $\mathcal{J}_F$ acts as a mapping from spatial frequency to the strength of presence of that frequency in the image.

`Python commands: --weighted_fourier: calculate frequency-weighted mean fourier coefficients; --show_fourier: display visualization`

***Discrete Wavelet Transform (DWT)***

This measure is based on how much information is contained in the largest DWT detail coefficients *D* extracted from an image *I* of size *M* x *N*.

$$S_{DWT} = \frac{1}{MN} \sum_{d_i \in D} |d_i|$$

By default, we define *D* as the set of the largest (in absolute value) 1% horizontal, vertical, and diagonal coefficients. 4 levels of DWT are applied using the Haar mother wavelet.

`Python commands: --wavelet_details: calculate discrete wavelet transform; --show_dwt: display visualization; --wt_threshold_percentile [#]: specifies the threshold percentile (default: 99) --wt_n_levels [#]: specifies number of decomposition levels (default: 4) --wt_mother_wavelet [wavelet type]: controls type of mother wavelet used  (default: haar)`

**(d) Generalized Variance (Heterogeneity)**

***Local (Intra-patch) Heterogeneity***

Estimates the mean local intra-patch covariance over the image, written:

$$\mathcal{C}_L(I) = \frac{1}{|P|} \sum_{p \in P} log\left(det\left(\hat{C}(p)\right) + 1\right),$$

where $\hat{C}(p)$ is the empirical covariance matrix of patch $p$.

Note that trace instead of determinant is used in the greyscale case.

```
Python commands: --local_covars: calculate local heterogeneity;
--show_local_covars: display visualization;
--local_cov_patch_size [#]: specifies patch size (default: 20);
--local_covar_wstep [#]: specifies distance between patches
(default: 5)
```

***Global (Inter-patch) Heterogeneity***

Computes the log-determinant of the global covariance matrix over patches in the image, where each multi-channel patch is unfolded into a single continuous vector, written:

$$\mathcal{C}_G(I) = log\left(det\left(\hat{C}(P)\right)\right),$$

where P is the set of vectorized patches, as in the case of Patch-wise Differential Entropy.

```
Python command: --global_patch_covar: calculate global
heterogeneity
```

##### (e) Pairwise Distance Measures (Patch Dissimilarity)

###### *Pairwise Distance Between Patch Means*

Computes the average pairwise distance between the first moments (means) of the patches across the image:

$$\mathcal{D}_{M,1}(I) = \frac{1}{|C||P|^2} \sum_{p_i, p_j \in P} ||\,\mu(p_i) - \mu(p_j)||_2$$

where $\mu(p) \in \mathbb{R}^3$ is the mean pixel value over patch *p*.

By default, this method utilizes *non-overlapping* patches (as do the remaining measures in this section).

`Python commands: --pairwise_mean_distances: calculate pairwise distance between patch means; --show_pw_mnt_ptchs: display patch visualization; --pw_mnt_dist_nonOL_WS [#,#]: specifies the number of patches when discretizing for the pairwise patch distance measures (default: 3,4)`

###### *Pairwise Distance Between Patch Moments*

This measure is similar to the one just above, except that it considers the second moment (the covariance) as well:

$$\mathcal{D}_{M,2}(I) = \frac{1}{|P|^2} \sum_{p_i, p_j \in P} \frac{1}{|C|} ||\,\mu(p_i) - \mu(p_j)||_2 + \frac{\gamma C}{|C|^2} ||\Sigma(p_i) - \Sigma(p_j)||_{1,1/2}$$

where $\sum(p) \in \mathbb{R}^{3\times3}$ is the covariance matrix of pixel values over the patch $p$ and $||M||_{1,1/2} = \sqrt{\sum_{k,\ell} |M_{k,\ell}|}$ is the simple entry-wise matrix norm.

By default, this method utilizes non-overlapping patches as well and sets $\gamma_C = \gamma_\mu = 1$.

```
Python commands: --pairwise_moment_distances: calculate pairwise distance between patch moments; --gamma_mu_weight [#]: specifies the weight on the mu distance (default: 1.0) --gamma_cov_weight [#]: specifies the weight on the covariance matrix distance (default: 1.0); --show_pw_mnt_ptchs: display patch visualization; --pw_mnt_dist_nonOL_WS [#,#]: specifies the number of patches when discretizing for the pairwise patch distance measures (default: 3,4)
```

***Distributional Divergences***

Since trivial vector space metrics (e.g., the $L_2$ distance) can be poor image metrics in the context of biological visual perception (e.g., Zhang et al. 2018), we consider the use of distributional divergences, which are more robust to perceptually irrelevant perturbations, such as small translations. We assume that our image has been separated into patches, and we have fit a simple empirical distribution to each one. We then compute complexity measures based on probabilistic divergences between these patch-wise distributions. Consider the case of using only the first two moments to approximate the distribution of a patch (i.e., using the empirical mean $\mu(p) \epsilon \mathbb{R}^3$ and covariance $\sum(p) \epsilon \mathbb{R}^{3\times3}$ for each patch $p$.). Then, following the natural Maximum Entropy assumption (Jaynes, 1957), we consider each patch to be multivariate Gaussian $\mathcal{G} = \mathcal{N}(\mu)(p), \sum(p)$, and assume that a complex image is likely to contain patches that are as "different" from each other as possible. The choice of a Gaussian distribution (1) renders many divergences readily computable and (2) assumes a minimally informative prior, given the two moments (i.e., it makes minimal statistical assumptions) (Jaynes, 1957). We therefore measure the expected pairwise distance between patches as a simple complexity measure.

We consider a family of complexity measures on an image $I$ defined by

$$\mathcal{C}_{\mathcal{D}}(I) = \frac{1}{|P|^2} \sum_{p_i, p_j \in P} \mathfrak{D}[\mathcal{G}_i \,||\, \mathcal{G}_j]$$

where $P$ is the set of patches, $\mathcal{G}_i$ is the Gaussian corresponding to patch $i$, and $\mathfrak{D}$ is a particular parameterizing choice of information-theoretic divergence, the options for which are detailed below. Note that these divergences (and hence the resulting complexity measures) are simple to compute and computationally tractable, due to the Gaussian assumption.

Implementation-wise, we consider $\mathfrak{D} \in \{\mathfrak{D}_J \mathfrak{D}_{W_2} \mathfrak{D}_B \mathfrak{D}_H \mathfrak{D}_{FMATF}\}$.

By default, this method utilizes *non-overlapping* patches as well and sets $\gamma_C = \gamma_\mu = 1$.

In the following, let $\mathcal{P}, \mathcal{Q}$ be probability distributions over $x \in X$ and $f_p, f_q$ be their respective densities. $X$ is the set of (continuous) pixel values, and our (per-patch) distributions $\mathcal{P}, \mathcal{Q}$ are simply fitted with the first two moments per patch. $d_\mu$ and $d_\sigma$ are simply two different "distances" for the means and the covariances, respectively. $\mu_i$ and $\mu_j$ are the means for $\mathcal{G}_i$ and $\mathcal{G}_i$, respectively.

Users can choose between Jeffrey's (symmetric KL) divergence (Jeffreys, 1946), the Wasserstein-2 metric (Kantorovich, 1939; Vaserstein, 1969), the squared Hellinger distance (Hellinger.1909), the Bhattacharyya distance (Bhattacharyya, 1943), and the Forstner-Moonen Abou-Moustafa-Torres-Ferries (FM-ATF) density metric (Förstner and Moonen 2003).

***Jeffrey's Divergence***

Simply the symmetric KL-divergence:

$$\mathfrak{D}_J[\mathcal{G}_i \mid\mid \mathcal{G}_j] = \frac{1}{2}\mathfrak{D}_{KL}[\mathcal{G}_i \mid\mid \mathcal{G}_j] + \frac{1}{2}\mathfrak{D}_{KL}[\mathcal{G}_j \mid\mid \mathcal{G}_i]$$

where

$$\mathfrak{D}_{KL}[\mathcal{P}, \mathcal{Q}] = \int_X f_p(x) \, log\left(\frac{f_p(x)}{f_q(x)}\right) dx$$

`Python commands: --pwg_jeffreys_div: calculate Jeffrey's divergence; --show_pw_mnt_ptchs: display patch visualization; --pw_mnt_dist_nonOL_WS [#,#]: specifies the number of patches when discretizing for the pairwise patch distance measures (default: 3,4)`

##### ***Wasserstein-2 Metric***

For the specific case of Gaussian distributions, the Wasserstein-2 distance simplifies down to the following form:

$$\mathfrak{D}_{W_2}[\mathcal{G}_i \,||\, \mathcal{G}_j] = ||\mu_i - \mu_j||\,_2^2 + tr\left(\Sigma_i + \Sigma_j - 2\left[\Sigma_j^{1/2}\Sigma_i\Sigma_j^{1/2}\right]^{1/2}\right)$$

```
Python commands: --pwg_w2_div: calculate Wasserstein-2 metric;
--show_pw_mnt_ptchs: display patch visualization;
--pw_mnt_dist_nonOL_WS [#,#]: specifies the number of patches
when discretizing for the pairwise patch distance measures
(default: 3,4)
```

##### ***Bhattacharyya Distance***

For notational brevity, we define the Bhattacharyya coefficient as:

$$\mathcal{B}[\mathcal{P},\mathcal{Q}] = \int_X \sqrt{f_p(x)f_q(x)}dx$$

Then the Bhattacharyya distance is simply given by:

$$\mathfrak{D}_B[\mathcal{G}_i \,||\, \mathcal{G}_j] = -log\mathcal{B}[\mathcal{G}_j,\mathcal{G}_i]$$

```
Python commands: --pwg_bhattacharyya_div: calculate
Bhattacharyya distance;
--show_pw_mnt_ptchs: display patch visualization;
```

--pw_mnt_dist_nonOL_WS [#,#]: specifies the number of patches when discretizing for the pairwise patch distance measures (default: 3,4)

***Squared Hellinger Distance***

Utilizing the notation just above, the squared Hellinger distance is given by:

$$\boldsymbol{\mathfrak{D}_H[\mathcal{G}_i \,||\, \mathcal{G}_j] = \sqrt{1 - \mathcal{B}[\mathcal{G}_i, \mathcal{G}_j]}}$$

Python commands: --pwg_hellinger_div: calculate squared Hellinger distance;

--show_pw_mnt_ptchs: display patch visualization;

--pw_mnt_dist_nonOL_WS [#,#]: specifies the number of patches when discretizing for the pairwise patch distance measures (default: 3,4)

***Forstner-Moonen Abou-Moustafa-Torres-Ferries (FM-ATF) Density Metric***

Folllowing Abou–Moustafa et al. 2010 and Förstner et al. 2003 we utilize the following metric:

$$\mathfrak{D}_{FMATF}[\mathcal{G}_i \,||\, \mathcal{G}_j] = d_\mu(\mathcal{G}_i, \mathcal{G}_j)^{1/2} + d_\sigma(\mathcal{G}_i, \mathcal{G}_j)^{1/2}$$

where the normalized (Mahalanobis) distance between means is given by:

$$d_\mu(\mathcal{G}_i, \mathcal{G}_j) = (\mu_i - \mu_j)^T \Sigma_a^{-1} (\mu_i - \mu_j)$$

with $\Sigma_a = (\Sigma_i + \Sigma_j)/2$ and the squared FM metric (Förstner et al. 2003) on Symmetric Positive Definite (SPD) matrices is written:

$$d_\sigma(\mathcal{G}_i, \mathcal{G}_j) = \sum_{\ell} log(\lambda_\ell)^2$$

with $diag(\lambda_1, \ldots, \lambda_K) = \Lambda$ satisfying the generalized eigenvalue problem $\Sigma_i V = \Lambda \Sigma_j V$.

```
Python commands: --pwg_fmatf_div: calculate FM-ATF density
metric; --show_pw_mnt_ptchs: display patch visualization;
--pw_mnt_dist_nonOL_WS [#,#]: specifies the number of patches
when discretizing for the pairwise patch distance measures
(default: 3,4)
```

We remind the reader that all the distributional divergences (i.e., in this sub-section) are computed using the patch-wise Gaussians.

#### S4: SUPPLEMENTAL DESCRIPTION OF DISCRETE WAVELET TRANSFORM (DWT) OUTPUT IMAGE

For a discrete wavelet transform, the signal is passed through a series of low-pass and high-pass filters. The high-pass filters will pass high frequency components (i.e., detail coefficients) and reject low frequency components (i.e., approximation coefficients) and vice-versa. The wavelet transform of an image produces four output images per decomposition level: an approximation image and a horizontal, vertical, and diagonal detail image. See schematic of discrete wavelet transform in Figure S2.

- The approximation image is obtained by vertical and horizontal low-pass filtering (i.e., only contains low frequency content, or approximation coefficients)
- The horizontal detail image is obtained by vertical high-pass and horizontal low-pass filtering
- The vertical detail image is obtained by vertical low-pass and horizontal high-pass filtering
- The diagonal detail image is obtained by vertical and horizontal high-pass filtering

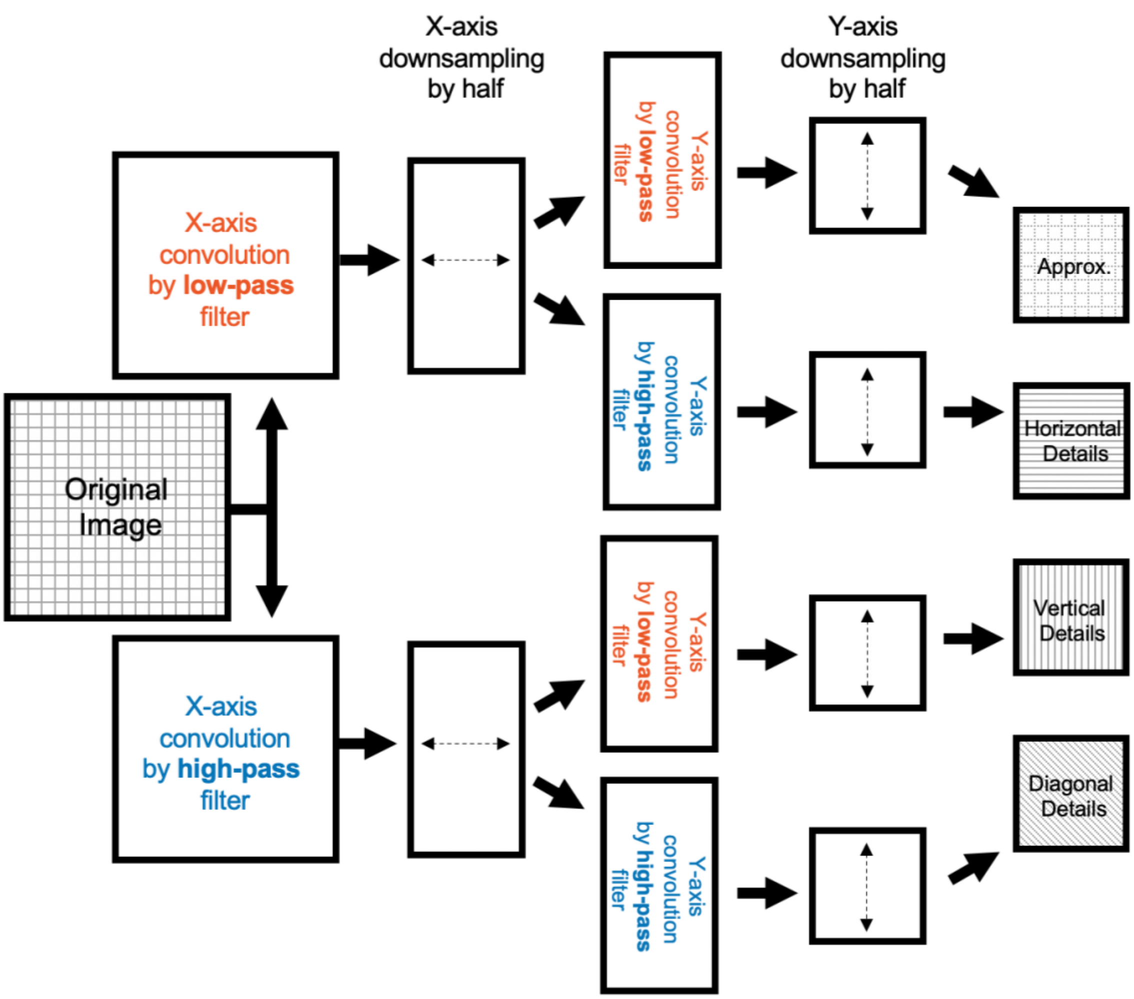


**Figure S2.** Schematic of discrete wavelet transform algorithm for an image.

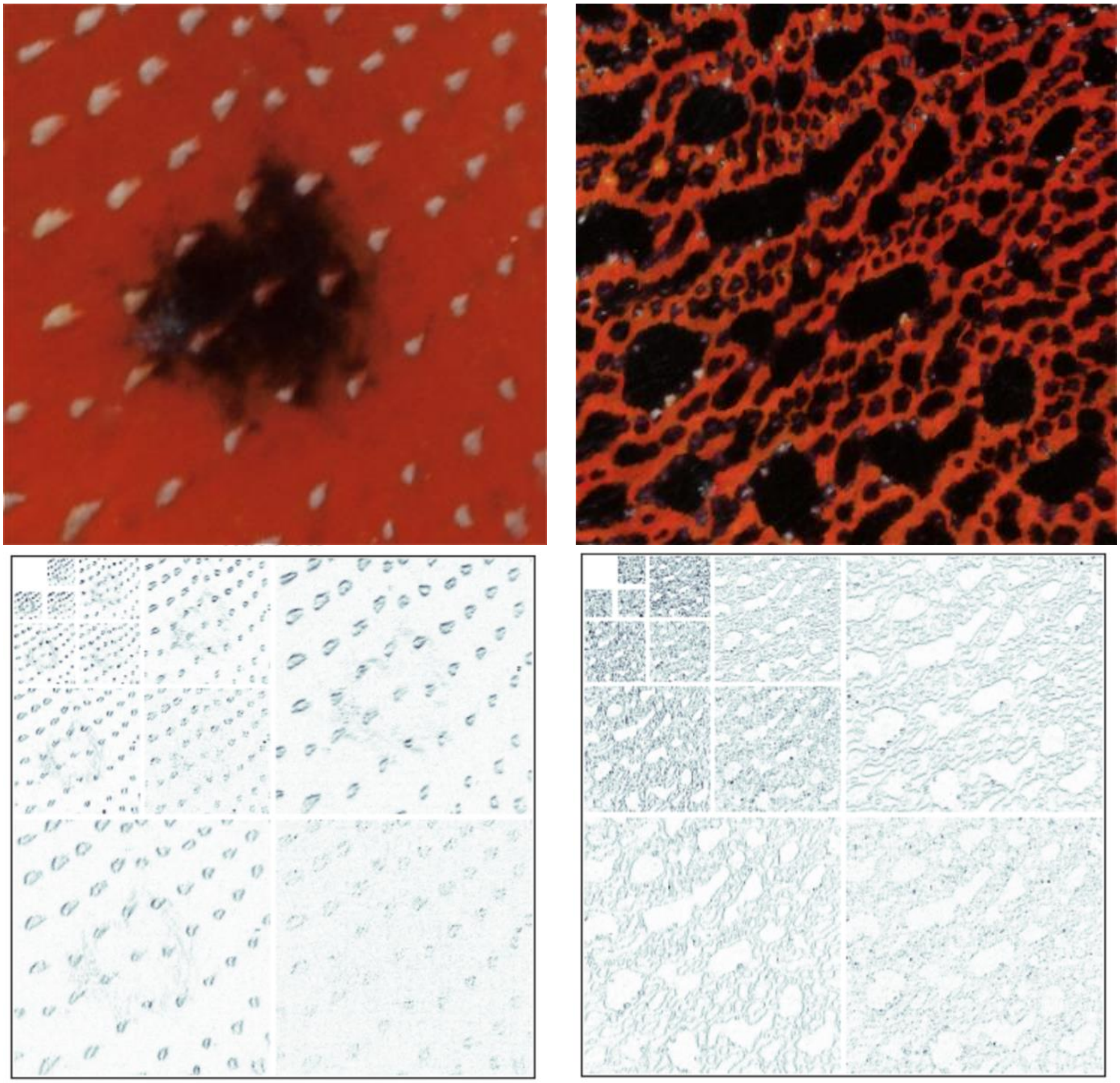

**Figure S3**. Output image for 4-level discrete wavelet transform for selected regions of digital photographs of two male *Anolis* lizard dewlap throat fans (*Anolis lyra on left; A. antioquiae on right).* The top panels contains the original images. In each bottom panel, the detail images decrease in size per decomposition level, with the largest subpanels being detail images of decomposition level 1, and the smallest being the detail images of decomposition level 4. For each decomposition level, the top right image represents the horizontal details, the bottom right image represents the diagonal details, and the bottom left image represents the vertical details.

**S5: SUPPLEMENTAL IMAGE SEGMENTATION DESCRIPTION AND EXAMPLE**

DEWPAT uses the scikit-learn package (Pedregosa et al. 2011) to implement three major clustering options: density-based spatial clustering of applications with noise (DBSCAN; Ester et al. 1996), graph cut optimization (graph-cuts; Greig, Porteous, and Seheult, 1989), and k-means clustering (Lloyd, 1982). We briefly describe these methods in the main text and have provided a more detailed overview in Table S2. In figure S4, we provide an example to illustrate the different potential outputs of each method by segmenting the elytra in an image of the beetle *Sternotomis pulchra* (taken from the online Lamiinae gallery; Roguet, 2022). Figure S4 shows the resulting output with original image used for each segmentation (shown only in the k-means panel for clarity), the clusters identified (coloured to show maximum differences), the clusters overlaid onto the original image, and the resulting segmented image. For k-means clustering, DEWPAT can return images coloured by the mean, median, or mode pixel colour for each cluster. It should be noted that DBSCAN and graph-cuts segment images based on pixel spatial proximity and colour similarity. Obtaining desired output relies on the user carefully selecting appropriate parameters. The diverse range of parameter choices and combinations make DBSCAN and graph-cuts best suited for cases in which images contain simple patterns and are similar to one another. As k-means clustering only considers colour similarity, the main parameter to consider is the number of clusters (K). As K can be chosen per image or automatically assigned, k-means clustering is well suited for cases in which images contain complex patterns and are different from one another. We provide details on the main arguments used in seg.py in Table S3.

**Table S2.** General overview of the differences between clustering methods.

| | **DBSCAN** | **Graph-cuts** | **K-means** |
|---|---|---|---|
| **Algorithm type** | Density-based clustering algorithm | Graph optimization algorithm | Centroid-based clustering algorithm |
| **Clustering description** | Groups together points that are closely packed together based on a distance measure | Image is represented as a graph in which nodes represent pixels and edges represent relationships between pixels. Graph-cuts optimizes an energy function over the graph, partitioning the graph into segments that minimize the energy | Partitions data into K clusters in which each data point belongs to the cluster with the nearest mean |
| **Pixel features considered** | Spatial proximity and colour similarity | Spatial proximity and colour similarity | Colour similarity only |
| **Parameter definition** | `--dbscan_eps`: epsilon (ε), the radius within which the algorithm looks for neighboring points<br><br>`--dbscan_min_neb_size`: minimum number of points required to form a dense region | `--gc_compactness`: controls the compactness of the superpixels used in the graph construction<br><br>`--gc_n_segments`: specifies the initial number of superpixels used to initialize the segmentation<br><br>`--gc_slic_sigma`: determines the scale or spread of the Gaussian kernel used in superpixel initialization, influencing the spatial extent over which pixel similarities are computed | `--kmeans_k`: specifies number of clusters<br><br>`--kmeans_k_file_list`: specifies path to .csv file that lists k values per image<br><br>`--kmeans_auto_crit`: criterion for choosing the best k-value for automatic k-means clustering (silhouette, davies_bouldin, calinski_harabasz)<br><br>`--kmeans_auto_bounds`: upper and lower bounds for cluster number |
| **Parameter impact** | dbscan_eps:<br>↑ values: fewer, larger clusters, with points within each cluster being more spatially dispersed (if set too high, may merge distinct clusters)<br>↓ values: more, smaller clusters, with points within each cluster being more spatially concentrated (if too low, points may be considered noise and not cluster)<br><br>dbscan_min_neb_size:<br>↑ values: fewer, more compact clusters with higher confidence in cluster membership (if set too high, may merge distinct clusters)<br>↓ values: more, potentially overlapping clusters (if too low, noise may cluster) | gc_compactness:<br>↑ values: encourage superpixels to adhere more closely to colour and spatial homogeneity leading to smoother boundaries but may result in over smoothing/loss of detail<br>↓ values: may result in more detailed but potentially less coherent segments<br><br>gc_n_segments:<br>↑ values: finer segmentation with more detail<br>↓ values: fewer but larger segments<br><br>gc_slic_sigma:<br>↑ values: smoother and more coherent superpixels<br>↓ values: captures finer details but may lead to over-segmentation | kmeans_k: all images will be segmented to same number of clusters<br><br>kmeans_k_file_list: each image can be segmented into different number of clusters<br><br>kmeans_auto_crit: different criteria change automatic clustering result<br><br>kmeans_auto_bounds: automatic clustering will only consider values within specified bounds |
| **Parameter defaults** | dbscan_eps: 1<br><br>dbscan_min_neb_size: 20 | gc_compactness: 10<br><br>gc_n_segments: 300<br><br>gc_slic_sigma: 0 | kmeans_k: chosen automatically<br><br>kmeans_k_file_list: None<br><br>kmeans_auto_crit: Davies-Bouldin criteria<br><br>kmeans_auto_bounds: 2, 6 |
| **Best suited for** | Images with simpler, more similar colour patterns | Images with simpler, more similar colour patterns | Images with complex, diverse patterns |

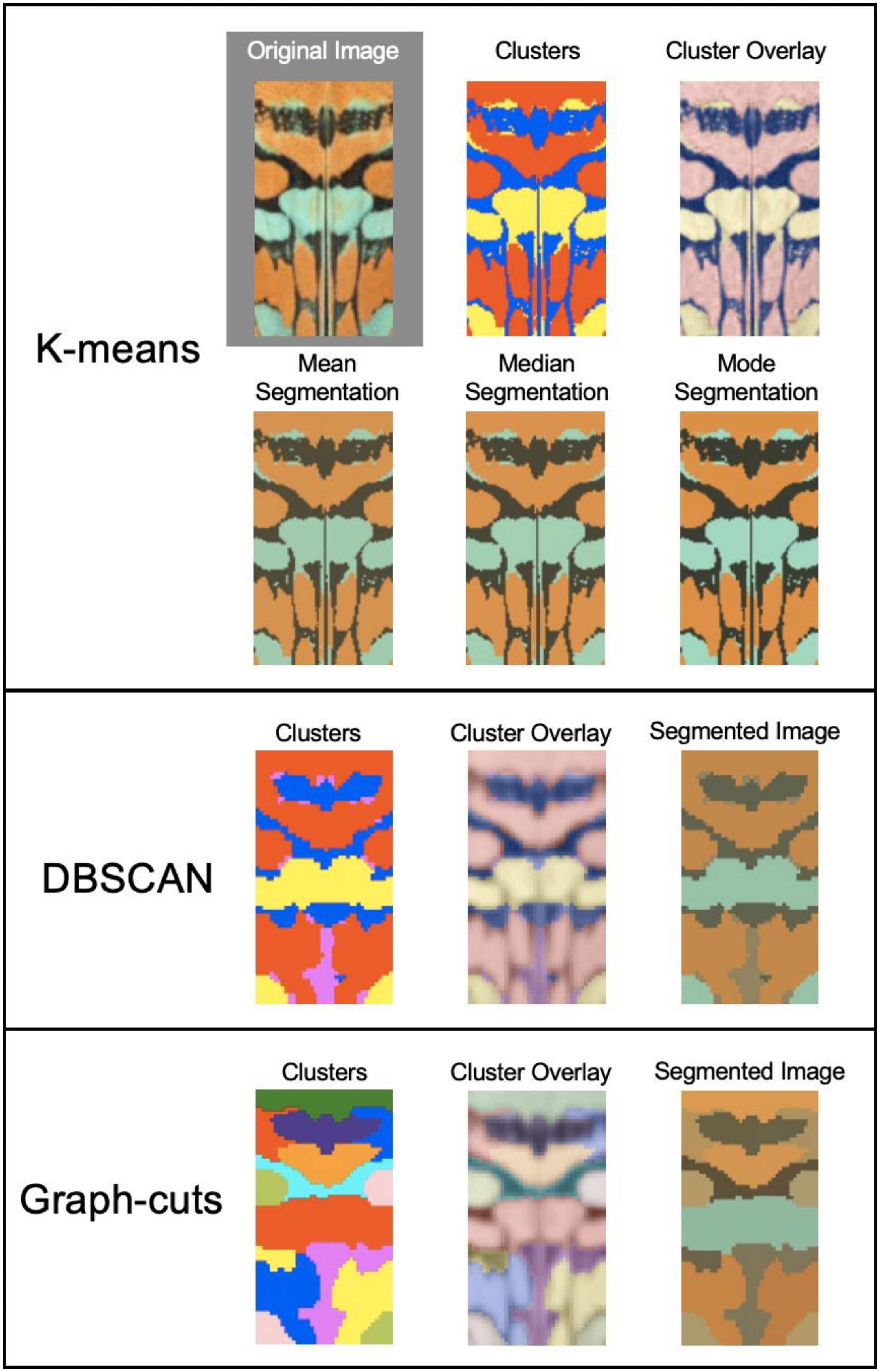
K-means
Original Image
Clusters
Cluster Overlay
Mean Segmentation
Median Segmentation
Mode Segmentation
DBSCAN
Clusters
Cluster Overlay
Segmented Image
Graph-cuts
Clusters
Cluster Overlay
Segmented Image

**Figure S4.** *Sternotomis pulchra* elytra segmented with k-means (top panel), DBSCAN (middle panel), and graph-cuts (bottom panel). The original image used in all segmentations is in the top right of the k-means panel. Panels show the identified clusters, those clusters overlaid onto the original image, and the resulting segmentation.

**Table S3.** Summary of main arguments used to run image segmentation on images in seg.py `DEWPAT` script.

| | DBSCAN | Graph-cuts | K-means |
|---|---|---|---|
| **General** | `--h` or `--help`: display help page; `--display`: display visualization (e.g., Figure S4); `--no_print_transitions`: prevent printing the transition matrix (prints by default), speeds up computation time; `--rseed [#]`: specify a fixed random seed to use for this run; `--clustering_colour_space [rgb,hsv,cie,lab]`: specify the colour space in which to perform the clustering (default: rgb) | | |
| **Specify clustering algorithm** | `--labeller dbscan`: use dbscan algorithm | `--labeller graph_cuts`: use graph-cuts algorithm | `--labeller kmeans`: use kmeans algorithm |
| **Labeller parameters** | `--dbscan_eps [#]`: epsilon neighbourhood (default: 1.0)<br><br>`--dbscan_min_neb_size [#]`: min samples needed (default: 20) | `--gc_compactness [#]`: superpixel compactness. (default: 10.0)<br><br>`--gc_n_segments [#]`: number of segments used in initialization (default: 300)<br><br>`--gc_slic_sigma [#]`: Gaussian kernel width (default: 0.0) | `--kmeans_k [#]`: specify number of clusters (chosen automatically by default)<br><br>`--kmeans_k_file_list [.csv file]`: specifies path to csv file that lists k values per image (e.g., "image.png,5")<br><br>`--kmeans_auto_bounds [#,#]`: lower and upper bounds on the number of clusters searched over for kmeans (default: 2,6)<br><br>`--kmeans_auto_crit [silhouette,davies_bouldin,calinski_harabasz]`: choice of criterion for choosing the best k-value (default: davies_bouldin) |
| **Small cluster merging parameters** | NA | NA | Sometimes automatic kmeans clustering algorithm creates clusters with very few pixels. To combat this, we've included a few options to merge small clusters into closest larger cluster.<br><br>`--merge_small_clusters_method [k_dependent,fixed]`: specify how to merge small clusters (default: none)<br><br>`--fixed_cluster_size_merging_threshold [#]`: fixed threshold percentage for cluster merging (only used when merging method is fixed) (default: 0.1)<br><br>`--small_cluster_merging_kdep_param [#]`: parameter for k_dependent annealed thresholding (initial thresh for k=2) (default: 0.05)<br><br>`--small_cluster_merging_dynamic_k`: if using k_dependent merging threshold, specify to recompute the threshold after every merge (default: False) |
| **Output segmented images** | `--display`: display visualization | `--display`: display visualization | The following arguments (1) tell `DEWPAT` to save segmented images and (2) specify the folder to which saved segmented images must be written for cluster-mean, cluster-median, or cluster-mode. Both arguments and folder name must be specified.<br><br>`--write_mean_segs --mean_seg_output_dir [folder]`<br><br>`--write_median_segs --median_seg_output_dir [folder]`<br><br>`--write_mode_segs --mode_seg_output_dir [folder]` |
| **Output colour statistics** | NA | NA | The following arguments specifiy output file to which segment statistics with cluster-mean, cluster-median, or cluster-mode values should be written<br><br>`--seg_mean_stats_output_file [file.csv]`<br><br>`--seg_median_stats_output_file [file.csv]`<br><br>`--seg_mode_stats_output_file [file.csv]` |

| | | | |
|---|---|---|---|
| | | | `--cluster_number_file [file.csv]`: specifies output file to which the number of estimated clusters (found with automatic kmeans clustering) per image should be written |
| **Example code used in figure S4** | `python seg.py Sternotomini.png --labeller dbscan --dbscan_eps 10 --dbscan_min_neb_size 35 --display` | `python seg.py Sternotomini.png --labeller graph_cuts --gc_compactness 30 --gc_n_segments 30 --gc_slic_sigma 0 --display` | `python seg.py Sternotomini.png --labeller kmeans --display` |

#### S6: SUPPLEMENTAL DATA OUTPUT AND VISUALIZATION OPTIONS

DEWPAT provides quantitative numerical output and includes qualitive visualization capabilities for understanding how colour and pattern are distributed within images. After running pattern analysis tools, users can output a .csv file containing numerical complexity scores for each image using `>file_name.csv`. By default, DEWPAT will run and output all complexity measures. Use arguments found above or in the package help page (`python img_complexity.py --help`) to specify specific measures. We note that while each DEWPAT tool is focused on measuring a particular axis of visual pattern complexity, many of these axes may exhibit interesting or informative correlations in empirical systems. Users may wish to study the dimensionality of the pattern complexity attributes measured with DEWPAT using ordination methods, such as a principal components analysis (PCA). For standard RGB images, users can specify whether to display a visual representation of extracted pattern information for each major measure of complexity (information entropy, edge content, detail granularity, heterogeneity, and patch dissimilarity; use `--show_all` to see all visualizations).

Images transformed with AcuityView (Caves & Johnsen, 2017; `preprocess.py`) or via colour segmentation (`seg.py`) are output to a specified folder. To use AcuityView (Caves & Johnsen, 2017), run:

```
python preprocess.py [input] [csv] [output]
```

Where input is an image (e.g., image.png) or a folder of images (e.g., folder_name), csv is a csv file containing image name (including file extension), width, viewing distance, and MRA (minimum resolvable angle) for each image, and output is a folder to store the blurred images.

The segmentation feature also provides quantitative outputs including a .csv file containing the colour code (in either RGB, HSV, or CIELAB colour space) and pixel frequency of each cluster for each image, and, if automatic clustering is performed, a .csv file with the number of clusters detected for each image. Additionally, `DEWPAT` is also capable of calculating a transition matrix for each segmented image (based on Endler's adjacency analysis; Endler, 2012; van der Berg et al., 2020). Briefly, this matrix consists of the transitions between colour classes taken from pairs of adjacent points along rows and columns of the image and provides information including relative area and proportion of colour classes (see Endler, 2012 for detailed description of methodology and uses). See table S3 for summary of image segmentation python commands.

For standard RGB images, `vis.py` includes the option to display a variety of plots summarizing pixel value distributions in RGB and HSV colour spaces (use `--all` to display all visualizations). One example is the 1D Colour Histogram feature (`--manual_unfolded_1d`), which displays a histogram of the pixel values with respect to a range of colour bins (in-text Figure 3). To visualize the standard three-channel (RGB) colour space in 1D and construct a histogram of image colours, we take the following trivial approach: (i) define a set of key points in RGB space, (ii) construct a 1D binning based on the ordering of (i), and (iii) associate each image pixel to a 1D bin, to form the histogram. For (i), one can simply choose a set of colours to act as "key points", with linear interpolation of colours applied between key points. Alternatively, any standard colourmap (function from [0,1] to RGB; e.g., from Matplotlib) can be used. Then, for (ii), we discretize the 1D space into bins, sampling the map to obtain a colour associated to each 1D bin, based on the interpolated key points or colourmap. Finally, for (iii), we bin the image pixels into 1D, via associating each pixel to the bin with the closest colour; the counts of this binning form the histogram. We remark that these maps, from the unit cube (here, 3D colours from sRGB images) to 1D, are effectively arbitrary (not wavelength based), and meant only for visualization. Users have the option to output these histogram values to a .csv file (`--write_1d_histo_vals --output_file [file.csv]`), which can be used to calculate colour diversity (e.g., using the Shannon-Weiner diversity index; Shannon & Weaver 1949).

601

602

603
